# Back to BERT in 2026: ModernGENA as a Strong, Efficient Baseline for DNA Foundation Models

**DOI:** 10.64898/2026.04.21.719816

**Authors:** Alena Aspidova, Yuri Kuratov, Artem Shadskiy, Mikhail Burtsev, Veniamin Fishman

## Abstract

Recent advances in DNA language models have mainly come from building larger and more complex architectures, making it harder to understand the effect of changes to standard components such as the transformer layers widely used in NLP. In this work, we study whether and how a modernized BERT-style backbone (ModernBERT) can be adapted to genomic sequence modeling to improve computational efficiency, training stability, and downstream performance. Under controlled experimental settings, we benchmark efficiency across a range of sequence lengths and evaluate downstream performance on the Nucleotide Transformer benchmark. The resulting model, ModernGENA, achieves a strong efficiency–quality trade-off and ranks among the top-performing models in our evaluation suite. To support reproducibility and provide a solid default reference point for future architectural work in genomics, we release the full implementation and configuration of ModernGENA as an open, reusable baseline, and make ModernGENA base and ModernGENA large publicly available through [the DNA language models collection on Hugging Face](https://huggingface.co/collections/AIRI-Institute/dna-language-models).

## 1 Introduction

DNA encodes a vast amount of biologically meaningful information, including regulatory logic, evolutionary constraints, and functional signals, that we are still far from fully decoding. Deep learning offers a promising route to learn general-purpose representations directly from sequence.

Existing approaches in genomics broadly follow two paradigms. Task-specific supervised models are trained end-to-end for particular tasks, such as Enformer (Avsec et al., 2021), Borzoi (Linder et al., 2025) and AlphaGenome (Avsec et al., 2026). In contrast, DNA foundation models first learn general-purpose sequence representations via self-supervised pretraining and are then adapted to downstream applications. Motivated by the goal of a single reusable backbone that transfers across many genomic tasks, DNA foundation modeling has expanded across architectural families, including CNN-based models such as ConvNova (Bo et al., 2025), Transformer models such as DNABERT (Ji et al., 2021), GENA (Fishman et al., 2025), DNABERT-2 (Zhou et al., 2023), and Nucleotide Transformer (Dalla-Torre et al., 2025), and SSM-inspired methods such as Caduceus (Schiff et al., 2024). Recent progress increasingly relies on composite, multi-module systems that integrate representations across scales, such as GENERator (Wu et al., 2025), Evo2 (Brixi et al., 2025), and Nucleotide Transformer v3 (Boshar et al., 2025). While often effective, these designs tend to increase computational cost by simultaneously scaling sequence length, representational resolution, and model size, thereby raising both the memory footprint and runtime requirements for training and inference.

Amid this emphasis on scaling and architectural complexity, it is easy to overlook continued progress in standard architectures. Recent Transformer refinements improve computational efficiency, training stability, and long-context handling, as in ModernBERT (Warner et al., 2025). Here we evaluate how well these improvements transfer to the genomics setting. This yields a strong baseline for systematic comparison of future advances and provides a modernized Transformer module that can be reused when building more efficient architectures, where Transformers often serve as a key component.

### Contributions

We evaluate how modern Transformer refinements transfer to genomics. Specifically, we

1. adapt and evaluate modern Transformer refinements for DNA foundation modeling, following ModernBERT advances;
2. benchmark efficiency–quality trade-offs under controlled experimental settings;
3. release training & finetuning code, configuration, and pretrained models ModernGENA base (135M) and ModernGENA large (377M) as a strong baseline for future work.

## 2 Related work

Among contemporary encoder-style foundation models for DNA, three widely used baselines are DNABERT-2 (Zhou et al., 2023), GENA-LM (Fishman et al., 2025), and Nucleotide Transformer (Dalla-Torre et al., 2025). While all follow masked-language pretraining (MLM), they differ most clearly in positional mechanisms, implementation of attention mechanism, and tokenization.

### Tokenization

NTv2 uses fixed k-mer tokens (6-mers), whereas DNABERT-2 and GENA-LM rely on variable-length BPE tokenization (Sennrich et al., 2016).

### Positional information

NTv2 incorporates rotary positional embeddings (RoPE) (Su et al., 2024) and expands its token-level context window; GENA-LM uses absolute positional encodings, while DNABERT-2 replaces explicit positional embeddings with ALiBi attention biases (Press et al., 2021) to reduce reliance on a learned position table and improve length extrapolation.

### Attention mechanism implementation

DNABERT-2 explicitly integrates FlashAttention (Dao et al., 2022) to speed up and reduce memory use in self-attention. While DNABERT-2 provides a FlashAttention-based implementation, we were unable to use it in our environment due to software incompatibilities: the installation procedure recommended in the official repository pins a Triton version that is incompatible with the released code. GENA-LM uses BERT-like full attention for shorter sequences and provides a specialized long-context variant with sparse attention (BigBird) (Zaheer et al., 2020). NTv2 retains full attention but modernizes the transformer block (including RoPE and more efficient MLP variants) to support a larger context window.

## 3 Experiments

### 3.1 Architecture

We research across two model sizes: **ModernGENA base** (135M parameters) and **ModernGENA large** (377M parameters). ModernBERT (Warner et al., 2025) serves as the backbone, which is an encoder-only Transformer modernized for stable training and high throughput on long sequences. The architecture incorporates the following key design choices:

- It scales better to long contexts in both representation quality and computational efficiency: it replaces absolute positional embeddings with RoPE (Su et al., 2024), and uses a hybrid attention pattern that alternates local sliding-window attention with global attention, with separate RoPE parameterizations for local and global layers.
- It improves optimization stability via a pre-norm block design (Xiong et al., 2020), an additional LayerNorm after the embedding layer, simplified normalization in the first attention block, and GeGLU in the FFN for a more expressive nonlinearity (Shazeer, 2020).
- It reduces unnecessary parameterization by disabling bias terms in most linear layers and in LayerNorm (except for the final linear layer).
- It accelerates training and inference through end-to-end unpadding (Zeng et al., 2022), variable-length kernel implementations (FlashAttention v3 (Shah et al., 2024) for global layers and v2 (Dao, 2023) for local layers), and compilation of compatible modules with torch.compile (Ansel et al., 2024).

Further details on the model architecture are provided in Appendix A.

### 3.2 Training

#### 3.2.1 Genomic datasets and preprocessing

The training corpus comprises all vertebrate species with available genome and transcript annotations in the NCBI RefSeq database as of December 2024 (443 assemblies), totaling 353,574,093,776 bp; the full list of assemblies is provided in Appendix F.

Previous studies have shown that DNA language models learn the structure of repetitive elements well, but perform worse on protein-coding and regulatory regions (Tang et al., 2025; Benegas et al., 2025b; 2023). To mitigate this bias, we follow a previously proposed sampling strategy (Brixi et al., 2025; Benegas et al., 2025a) and construct training intervals around gene starts, since these regions are depleted for simple repeats and enriched for regulatory signals and gene elements relevant to downstream genomic tasks. Specifically, for each gene and pseudogene, we extract a[−16 kbp, +8 kbp] window around each unique transcription start site. Overlapping windows are merged into non-overlapping regions using BEDTools. For each resulting region, we include both strands (forward and reverse complement). Sequences containing ambiguous nucleotides (symbols other than A/C/G/T) are excluded.

#### 3.2.2 Train/validation split

The data are partitioned at the level of whole chromosomes. For all assemblies except human (GCF 000001405.40), the validation set accounts for ∼10% of the total genome length, with the remaining ∼90% used for training. For the human assembly, chromosomes 8, 20, and 21 are held out for validation, and all remaining chromosomes are used for training.

#### 3.2.3 Tokenization

We use a 32k BPE vocabulary over the A/T/G/C/N symbols, with special tokens [CLS], [SEP], [PAD], [UNK], and [MASK] following (Fishman et al., 2025). As a preprocessing step, long runs of N are collapsed into a single token.

#### 3.2.4 Training settings

The model was trained on 8 NVIDIA A100 80GB GPUs with a global batch size of 4096 sequences. To improve robustness to variable input lengths, we used dynamic sequence packing with sequence lengths sampled uniformly from 10 to 1024 tokens (average length ≈ 700 tokens). We optimized using decoupled AdamW with learning rate 4 × 10^−4^, *β*_1_ = 0.9, *β*_2_ = 0.98, *ϵ* = 10^−6^, and weight decay 10^−5^. Weight decay was not applied to bias parameters or normalization layers. We used a warmup–stable schedule in token space: the learning rate was linearly warmed up over the first 3 × 10^9^ tokens and then held constant for the remainder of training.

ModernGENA base was trained for 158 epochs and processed 1,510B tokens in total. ModernGENA large was trained for 138 epochs, corresponding to 1,320B processed tokens. On 8× A100-80GB, the observed pretraining throughput was 1.33M tokens/s for ModernGENA base and 0.486M tokens/s for ModernGENA large.

### 3.3 Comparison group

To isolate the effect of modern Transformer-block refinements on performance and computational efficiency, the evaluation focuses on a *primary baseline set* of encoder-only Transformer models with fewer than 500M parameters: DNABERT-2, NTv2, and GENA-LM. Since NTv2 is available in multiple configurations spanning a wide range of parameter counts, NTv2 (100M) and NTv2 (250M) are selected to match the size of the ModernGENA models and ensure a fair comparison. When directly comparable metrics are available for other models with different architectures and model sizes, they are included as additional reference points, but they are not part of the primary baseline set.

### 3.4 Inference efficiency evaluation

Inference efficiency is measured as throughput (tokens/s) on fixed-length sequences using an NVIDIA A100 (80 GB) GPU. The evaluation considers the *primary baseline set* of encoder-only Transformer models. For each sequence length *L*, the maximum batch size that fits in memory is determined via exponential growth until an error, followed by binary search. Using this batch size, throughput is averaged over 10 timing runs per model (Appendix B).

Across these settings, ModernGENA achieves slightly higher throughput under the standard inference configuration and substantially higher throughput with FlashAttention 2, for both ModernGENA base and ModernGENA large (Figure 1).

**Figure 1:**
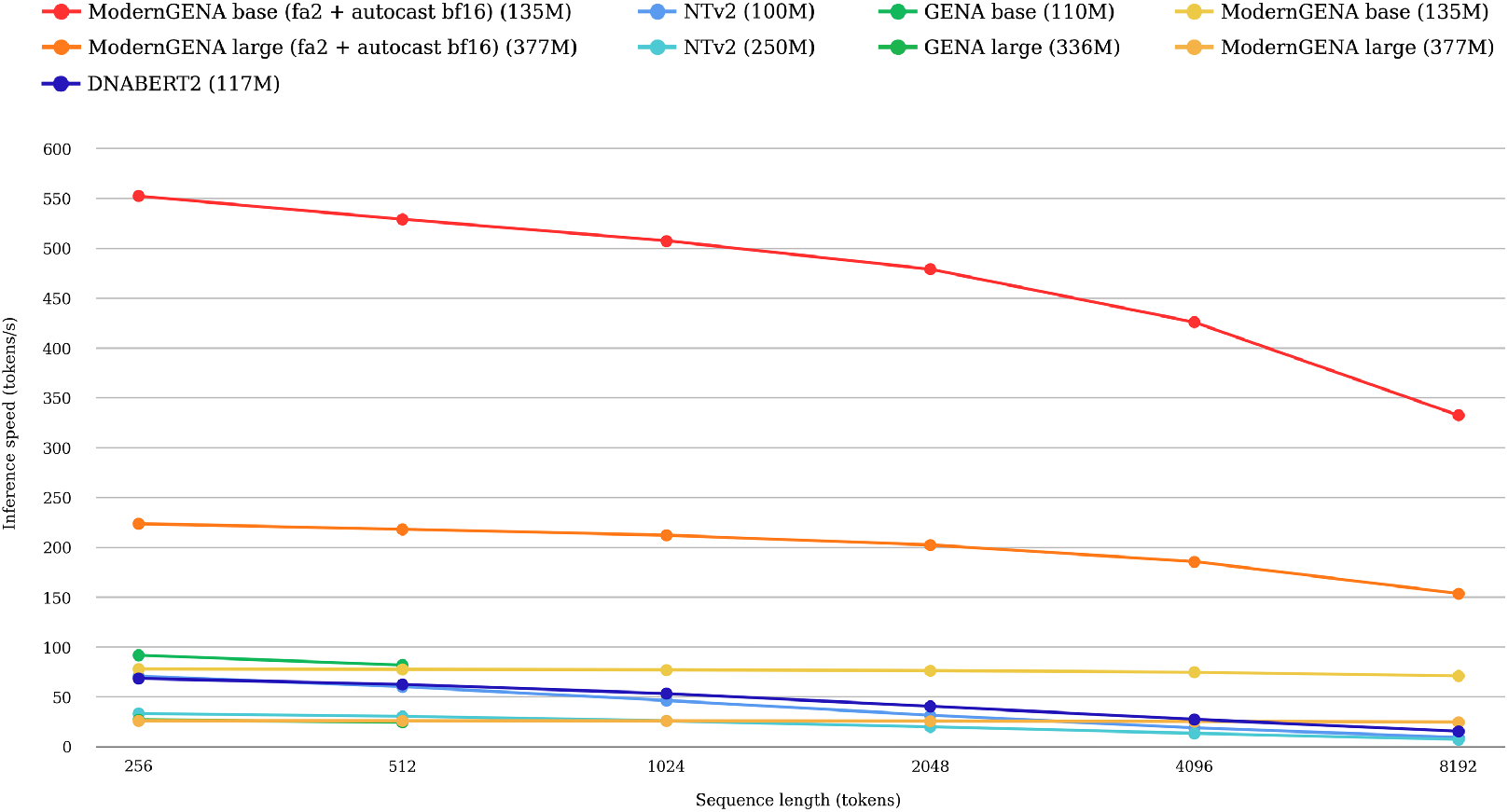
Inference efficiency on an NVIDIA A100 (80 GB). Models from the *primary baseline set* are benchmarked. Throughput is averaged over 10 timing runs per model.

### 3.5 Results on NT benchmark

To enable fair comparison with prior architectures, we evaluate ModernGENA on the NT benchmark (Dalla-Torre et al., 2025), which comprises 18 tasks. ModernGENA is fine-tuned separately for each task without reverse-complement (RC) augmentation or conjoining; full training details are provided in Appendix E. Performance is reported as Matthews correlation coefficient (MCC) under 10-fold cross-validation. Results for the remaining models are taken from (Dalla-Torre et al., 2025; Wu et al., 2025).

To summarize performance across all NT tasks, we rank models by MCC separately for each task and then average the resulting task-wise ranks. By this measure, ModernGENA shows competitive performance across a broad range of tasks, placing first within the *primary baseline set* and ModernGENA large second overall in the full comparison, behind GENERator (1.2B) (Figure 2). Full numerical results for both settings are provided in Appendix C and Appendix D.

**Figure 2:**
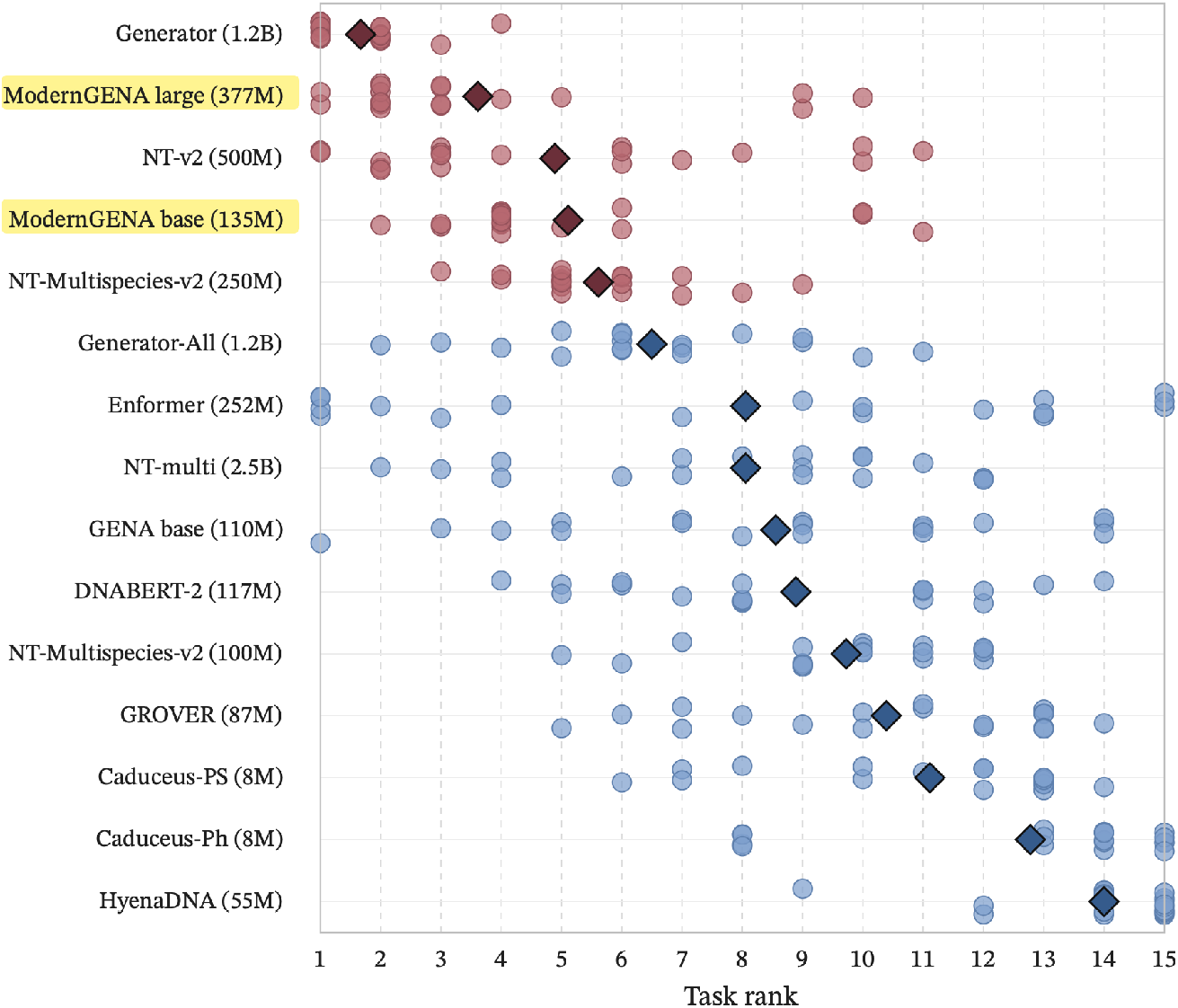
ModernGENA improves downstream performance and leads among comparable-size models. Circles denote task-specific ranks, while diamonds indicate the average rank across tasks.

We note that ModernGENA achieves lower scores than the strongest NT variants on the splice-site tasks. One plausible explanation is tokenization: splice prediction relies on short, precisely positioned sequence motifs, and fixed fine-grained tokenization may be better suited to capturing such signals than BPE.

This interpretation is consistent with recent findings that splice-site prediction particularly benefits from uniform, fine-grained tokenization (Kim et al., 2026). More broadly, this may point to the particular importance of tokenization choice, and of further work in this direction, in the genomics domain.

We further evaluated the usefulness of pretraining on the NT benchmark by focusing specifically on ModernGENA large and following the pretrained-versus-randomized comparison protocol introduced by Vishniakov et al.. Unlike the modest or architecture-dependent gains reported there for several genomic foundation models, ModernGENA large shows a consistent and substantial benefit from pretraining: average MCC improves from 0.420 with random initialization to 0.671 after pretraining, with improvements observed on all 18 NT tasks (Figure 3).

**Figure 3:**
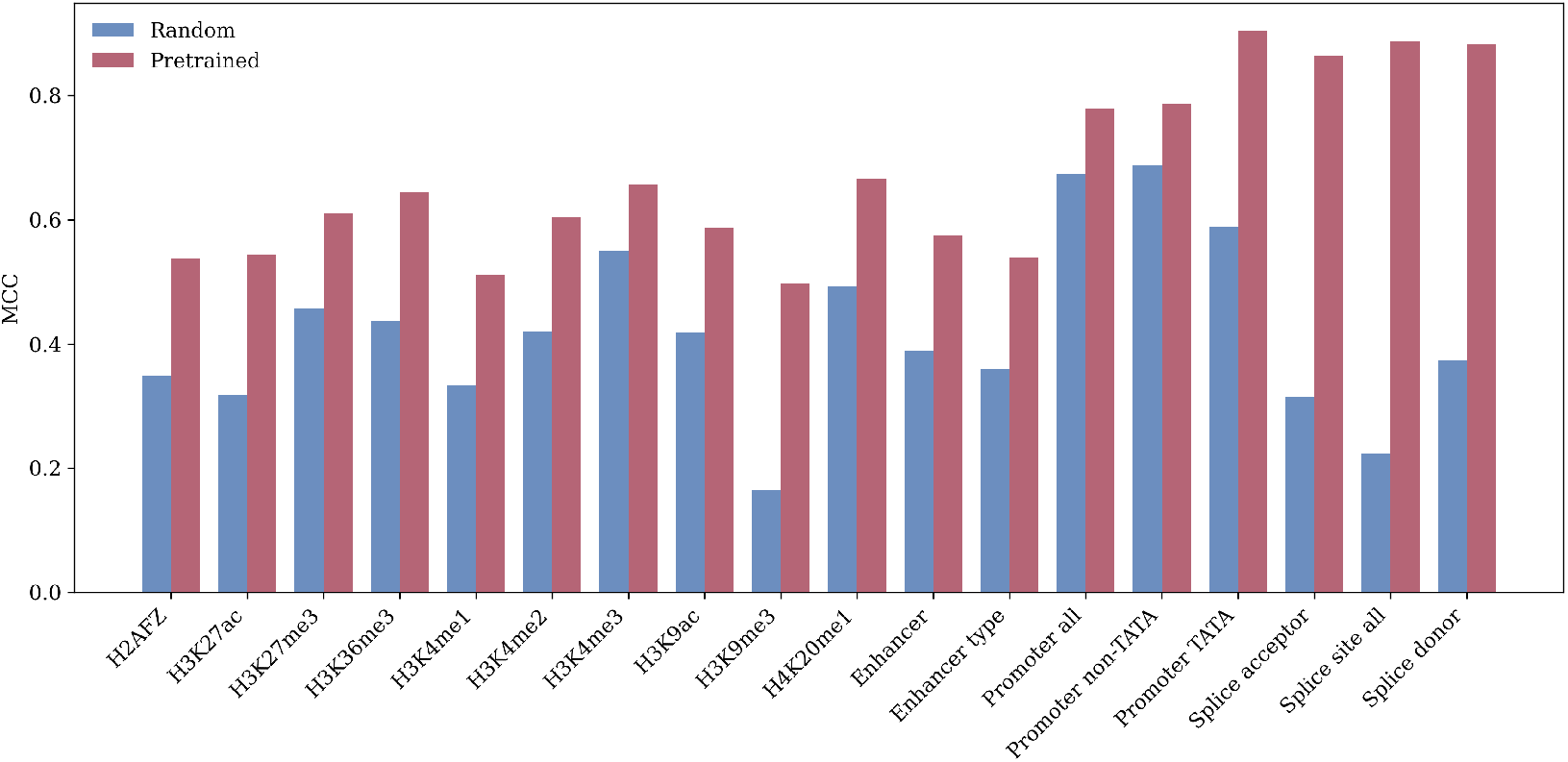
Masked-language pretraining consistently improves performance on wide range of genomic downstream tasks. Performance (MCC) is shown for each task comparing a randomly initialized model (random) and its pretrained counterpart.

## 4 Data availability

Pretrained ModernGENA models are publicly available on Hugging Face:

- moderngena-base
- moderngena-large

The training and fine-tuning code, including an example for fine-tuning ModernGENA, is available in the GENA LM repository at [github ModernGENA](https://github.com/AIRI-Institute/GENA_LM/tree/main/examples/modernGENA).

## 5 Conclusion

Standard Transformer encoders continue to improve through practical architectural and training modernizations that increase computational efficiency, robustness, and long-context capability. In this work, we study how these advances transfer to genomic sequence modeling by adapting ModernBERT for DNA pretraining and introducing ModernGENA.

We evaluate ModernGENA along two axes that matter in practice: efficiency and downstream transfer. Under controlled settings, we benchmark inference throughput across a range of sequence lengths and evaluate downstream performance on the Nucleotide Transformer benchmark. ModernGENA supports FlashAttention-based implementations and achieves higher inference throughput in our experiments, while delivering strong benchmark performance. ModernGENA ranks first among encoder-only models with comparable size and second overall in our evaluation suite, reflecting a favorable balance between efficiency and quality. Finally, because ModernGENA builds on the ModernBERT design, it is also intended to support long genomic contexts, a key requirement in many genomics applications.

Our results also point to a broader and still unresolved evaluation question in this area. Although the Nucleotide Transformer benchmark is one of the most widely used suites for comparing DNA foundation models, the absolute differences between architectures, and even between substantially different model scales, often appear modest. In practice, part of this gap can sometimes be reduced through careful tuning of fine-tuning hyperparameters and training protocols, which makes it difficult to draw strong conclusions from small metric differences alone. This raises an important open question: to what extent does increasing model scale translate into reliable downstream improvements for genomic sequence modeling, beyond what can be achieved with smaller models and strong training recipes? Answering this likely requires deeper analysis of benchmark characteristics and fine-tuning variance, and it motivates broader evaluation, either by incorporating additional benchmarks that stress different regimes (including longer-range context) or by reconsidering when scaling is actually necessary for genomic sequence understanding. In this context, we position ModernGENA as both a strong, reproducible baseline and a practical component for building and testing more complex architectures: it is efficient in throughput, adapted to long sequences, and intended to make future architectural experiments more systematic and comparable.

## Appendix A. Architecture configuration

Additional architecture configuration details for ModernGENA base and ModernGENA large are provided in Table 1.

**Table 1:** Architecture configuration.

| Parameter | ModernGENA base | ModernGENA large |
| --- | --- | --- |
| parameters | 135M | 377M |
| num_hidden_layers | 22 | 28 |
| hidden_size | 768 | 1024 |
| intermediate_size | 1152 | 2624 |
| num_attention_heads | 12 | 16 |
| attn_out_dropout_prob | 0.1 | 0.1 |
| mlp_layer | glu | glu |
| mlp_out_bias | false | false |
| activation_function | gelu | gelu |
| rotary_emb_base | 10000 | 10000 |
| sliding_window | 128 | 128 |
| global_attn_every_n_layers | 3 | 3 |

## Appendix B. Inference efficiency details

Inference efficiency is evaluated on an NVIDIA A100 (80 GB) using fixed-length sequences, reporting throughput (tokens/s) for models in the *primary baseline set*. The resulting batch sizes and tokens/s are reported in Table 2.

**Table 2:** Inference efficiency evaluation. For each sequence length *L*, the table reports the maximum batch size that fits in memory (bs) and throughput in tokens/s (mean over 10 runs).

| Model | Sequence length $L$ (tokens) | | | | | | | | | | | |
| --- | --- | --- | --- | --- | --- | --- | --- | --- | --- | --- | --- | --- |
|  | 256 |  | 512 |  | 1024 |  | 2048 |  | 4096 |  | 8192 |  |
|  | bs | tokens/s | bs | tokens/s | bs | tokens/s | bs | tokens/s | bs | tokens/s | bs | tokens/s |
| ModernGENA base (FA2 + autocast bf16) (135M) | 3631 | **552.44** | 1811 | **529.18** | 901 | **507.61** | 446 | **479.04** | 218 | **425.88** | 92 | **332.54** |
| ModernGENA base (135M) | 3631 | 78.12 | 1811 | 77.73 | 901 | 77.20 | 446 | 76.38 | 218 | 74.83 | 92 | 71.15 |
| GENA base (110M) | 6637 | 91.78 | 2206 | 81.98 | — | — | — | — | — | — | — | — |
| DNABERT-2 (117M) | 3802 | 68.57 | 1433 | 62.63 | 376 | 53.42 | 90 | 40.68 | 15 | 27.57 | 5 | 15.48 |
| NTv2 (100M) | 5941 | 70.88 | 1845 | 60.28 | 519 | 46.46 | 132 | 31.72 | 26 | 19.02 | 5 | 9.37 |
| ModernGENA large (FA2 + autocast bf16) (377M) | 2721 | **223.77** | 1356 | **218.23** | 673 | **212.24** | 332 | **202.51** | 161 | **185.82** | 67 | **153.77** |
| ModernGENA large (377M) | 2721 | 26.20 | 1356 | 26.11 | 673 | 26.01 | 332 | 25.85 | 161 | 25.56 | 67 | 24.86 |
| GENA large (336M) | 4922 | 27.13 | 1633 | 24.79 | — | — | — | — | — | — | — | — |
| NTv2 (250M) | 4107 | 33.47 | 1649 | 30.62 | 486 | 26.07 | 127 | 20.05 | 26 | 13.49 | 5 | 7.35 |

## Appendix C. NT benchmark results: primary baseline set

**Table 3:** NT Benchmark results (*primary baseline set*). 10-fold cross-validation, reported as 100 × MCC in the format *mean ± std* across folds. The final two rows report average ranks computed within the *primary baseline set* and within the full comparison across all models, respectively.

| Task | ModernGENA<br>base (135M) | ModernGENA<br>large (377M) | DNABERT-2<br>(117M) | GENA<br>base (110M) | NTv2<br>(100M) | NTv2<br>(250M) |
| --- | --- | --- | --- | --- | --- | --- |
| H2AFZ | 52.2 ± 0.7 | 53.7 ± 1.6 | 49.0 ± 1.3 | 49.5 ± 1.1 | 49.2 ± 1.2 | 51.3 ± 1.7 |
| H3K27ac | 51.2 ± 1.3 | 54.3 ± 1.0 | 49.1 ± 1.0 | 49.8 ± 1.7 | 48.7 ± 1.6 | 49.7 ± 1.4 |
| H3K27me3 | 60.8 ± 0.4 | 61.0 ± 0.8 | 59.9 ± 1.0 | 60.0 ± 0.7 | 59.5 ± 1.3 | 60.0 ± 0.9 |
| H3K36me3 | 63.8 ± 0.9 | 64.4 ± 1.0 | 63.7 ± 0.7 | 62.3 ± 0.9 | 61.7 ± 0.6 | 63.6 ± 1.2 |
| H3K4me1 | 49.3 ± 1.0 | 51.2 ± 0.9 | 49.0 ± 0.8 | 48.6 ± 0.9 | 48.5 ± 1.1 | 49.1 ± 0.6 |
| H3K4me2 | 57.5 ± 1.1 | 60.4 ± 1.3 | 55.8 ± 1.3 | 55.4 ± 0.8 | 55.1 ± 1.0 | 57.0 ± 0.9 |
| H3K4me3 | 64.9 ± 1.4 | 65.6 ± 2.0 | 64.6 ± 0.8 | 65.7 ± 1.2 | 63.3 ± 1.5 | 64.0 ± 0.9 |
| H3K9ac | 57.3 ± 0.8 | 58.6 ± 1.4 | 56.4 ± 1.3 | 54.3 ± 1.0 | 53.8 ± 1.5 | 56.5 ± 2.1 |
| H3K9me3 | 48.2 ± 1.8 | 49.7 ± 1.2 | 44.3 ± 2.5 | 49.1 ± 1.4 | 44.5 ± 1.7 | 46.7 ± 1.6 |
| H4K20me1 | 66.9 ± 0.6 | 66.6 ± 1.3 | 65.5 ± 1.1 | 65.6 ± 0.9 | 64.8 ± 0.8 | 65.2 ± 0.6 |
| Enhancer | 54.5 ± 0.6 | 57.5 ± 1.0 | 51.7 ± 1.1 | 53.6 ± 0.7 | 50.7 ± 0.9 | 52.5 ± 1.0 |
| Enhancer type | 51.2 ± 1.3 | 53.9 ± 0.7 | 47.6 ± 0.9 | 49.3 ± 0.7 | 46.5 ± 0.9 | 49.2 ± 1.0 |
| Promoter all | 77.4 ± 0.6 | 77.9 ± 0.7 | 75.4 ± 0.9 | 74.1 ± 1.1 | 75.3 ± 0.5 | 77.4 ± 1.3 |
| Promoter non-TATA | 77.9 ± 0.9 | 78.6 ± 0.4 | 76.9 ± 0.9 | 75.7 ± 0.8 | 76.6 ± 1.4 | 78.5 ± 1.0 |
| Promoter TATA | 90.6 ± 2.0 | 90.4 ± 1.2 | 78.4 ± 3.6 | 80.7 ± 4.9 | 82.6 ± 1.9 | 87.0 ± 1.9 |
| Splice acceptor | 85.1 ± 0.3 | 86.4 ± 0.5 | 83.7 ± 0.6 | 80.5 ± 0.9 | 94.7 ± 0.3 | 95.0 ± 0.8 |
| Splice site all | 87.6 ± 0.3 | 88.8 ± 0.7 | 85.5 ± 0.5 | 82.3 ± 0.8 | 96.0 ± 0.5 | 96.5 ± 0.3 |
| Splice donor | 86.8 ± 0.6 | 88.2 ± 0.8 | 86.1 ± 0.4 | 81.9 ± 0.7 | 94.7 ± 0.8 | 96.7 ± 0.4 |
| Average rank (primary baseline set) | **2.39** | **1.50** | **4.67** | **4.28** | **5.00** | **3.06** |
| Average rank (full comparison) | **5.11** | **3.61** | **8.89** | **8.56** | **9.72** | **5.61** |

## Appendix D. NT benchmark results: additional models not included in the primary baseline set

**Table 4:** NT Benchmark results including additional models not included in the primary baseline set. 10-fold cross-validation, reported as 100 × MCC in the format *mean ± std* across folds. The final row reports each model’s average rank across tasks.

| Task | ModernGENA<br>base (135M) | ModernGENA<br>large (377M) | NTv2<br>(500M) | NT-multi<br>(2.5B) | Enformer<br>(252M) | HyenaDNA<br>(55M) | Caduceus-Ph<br>(8M) | Caduceus-PS<br>(8M) | GROVER<br>(87M) | GENERator<br>(1.2B) | GENERator-All<br>(1.2B) |
| --- | --- | --- | --- | --- | --- | --- | --- | --- | --- | --- | --- |
| H2AFZ | 52.2 ± 0.7 | 53.7 ± 1.6 | 52.4 ± 0.8 | 50.3 ± 1.0 | 52.2 ± 1.9 | 45.5 ± 1.5 | 41.7 ± 1.6 | 50.1 ± 1.3 | 50.9 ± 1.3 | 52.9 ± 0.9 | 50.6 ± 1.9 |
| H3K27ac | 51.2 ± 1.3 | 54.3 ± 1.0 | 48.8 ± 1.3 | 48.1 ± 2.0 | 52.0 ± 1.5 | 42.3 ± 1.7 | 46.4 ± 1.8 | 46.4 ± 2.2 | 48.9 ± 2.3 | 54.6 ± 1.5 | 49.6 ± 1.4 |
| H3K27me3 | 60.8 ± 0.4 | 61.0 ± 0.8 | 61.0 ± 0.6 | 59.3 ± 1.6 | 55.2 ± 0.7 | 54.1 ± 1.8 | 54.7 ± 1.0 | 56.1 ± 3.6 | 60.0 ± 0.8 | 61.9 ± 0.8 | 59.0 ± 1.4 |
| H3K36me3 | 63.8 ± 0.9 | 64.4 ± 1.0 | 63.3 ± 1.5 | 63.5 ± 1.6 | 56.7 ± 1.7 | 54.3 ± 1.0 | 54.3 ± 0.9 | 60.2 ± 0.8 | 58.5 ± 0.8 | 65.0 ± 0.6 | 62.1 ± 1.3 |
| H3K4me1 | 49.3 ± 1.0 | 51.2 ± 0.9 | 49.0 ± 1.7 | 48.1 ± 1.2 | 50.4 ± 2.1 | 43.0 ± 1.4 | 41.1 ± 1.2 | 43.4 ± 3.0 | 46.8 ± 1.1 | 50.4 ± 1.0 | 49.0 ± 1.6 |
| H3K4me2 | 57.5 ± 1.1 | 60.4 ± 1.3 | 55.2 ± 1.3 | 55.2 ± 2.2 | 62.6 ± 1.5 | 52.1 ± 2.4 | 48.0 ± 1.3 | 52.6 ± 3.5 | 55.8 ± 1.2 | 60.7 ± 1.0 | 56.9 ± 1.2 |
| H3K4me3 | 64.9 ± 1.4 | 65.6 ± 2.0 | 62.7 ± 2.0 | 61.8 ± 1.5 | 63.5 ± 1.9 | 59.6 ± 1.5 | 58.8 ± 2.0 | 61.1 ± 1.5 | 63.4 ± 1.1 | 65.3 ± 0.8 | 62.8 ± 1.8 |
| H3K9ac | 57.3 ± 0.8 | 58.6 ± 1.4 | 55.1 ± 1.6 | 52.7 ± 1.7 | 59.3 ± 2.0 | 48.4 ± 2.2 | 51.4 ± 1.4 | 51.8 ± 1.8 | 53.1 ± 1.4 | 57.0 ± 1.7 | 55.6 ± 1.8 |
| H3K9me3 | 48.2 ± 1.8 | 49.7 ± 1.2 | 46.7 ± 4.4 | 44.7 ± 1.8 | 45.3 ± 1.6 | 37.5 ± 2.6 | 43.5 ± 1.9 | 45.5 ± 1.9 | 44.1 ± 1.7 | 50.9 ± 1.3 | 48.0 ± 3.7 |
| H4K20me1 | 66.9 ± 0.6 | 66.6 ± 1.3 | 65.4 ± 1.1 | 65.0 ± 1.4 | 60.6 ± 1.6 | 58.0 ± 0.9 | 57.2 ± 1.2 | 59.0 ± 2.0 | 63.4 ± 0.6 | 67.0 ± 0.6 | 65.2 ± 1.0 |
| Enhancer | 54.5 ± 0.6 | 57.5 ± 1.0 | 57.5 ± 2.3 | 52.7 ± 1.2 | 61.4 ± 1.0 | 47.5 ± 0.6 | 48.0 ± 0.8 | 49.0 ± 0.9 | 51.9 ± 0.9 | 59.4 ± 1.3 | 55.3 ± 2.0 |
| Enhancer type | 51.2 ± 1.3 | 53.9 ± 0.7 | 54.1 ± 1.3 | 48.4 ± 1.2 | 57.3 ± 1.3 | 44.1 ± 1.0 | 46.1 ± 0.9 | 45.9 ± 1.1 | 48.1 ± 0.9 | 54.7 ± 1.7 | 51.0 ± 2.2 |
| Promoter all | 77.4 ± 0.6 | 77.9 ± 0.7 | 78.0 ± 1.2 | 76.1 ± 0.9 | 74.5 ± 1.2 | 69.3 ± 1.6 | 70.7 ± 1.7 | 72.2 ± 1.4 | 72.1 ± 1.1 | 79.5 ± 0.5 | 76.5 ± 0.9 |
| Promoter non-TATA | 77.9 ± 0.9 | 78.6 ± 0.4 | 78.5 ± 0.9 | 77.3 ± 1.0 | 76.3 ± 1.2 | 72.3 ± 1.3 | 74.0 ± 1.2 | 74.6 ± 0.9 | 73.9 ± 1.8 | 80.1 ± 0.5 | 78.6 ± 0.7 |
| Promoter TATA | 90.6 ± 2.0 | 90.4 ± 1.2 | 91.9 ± 2.8 | 94.4 ± 1.6 | 79.3 ± 2.6 | 64.8 ± 4.4 | 86.8 ± 2.3 | 85.3 ± 3.4 | 89.1 ± 4.1 | 95.0 ± 0.9 | 86.2 ± 2.4 |
| Splice acceptor | 85.1 ± 0.3 | 86.4 ± 0.5 | 96.5 ± 0.4 | 95.8 ± 0.3 | 74.9 ± 0.7 | 81.5 ± 4.9 | 90.6 ± 1.5 | 93.9 ± 1.2 | 81.2 ± 1.2 | 96.4 ± 0.3 | 95.1 ± 0.6 |
| Splice site all | 87.6 ± 0.3 | 88.8 ± 0.7 | 96.8 ± 0.3 | 96.4 ± 0.3 | 73.9 ± 1.1 | 85.4 ± 5.3 | 94.1 ± 0.6 | 94.2 ± 1.2 | 84.9 ± 1.5 | 96.6 ± 0.3 | 95.9 ± 0.3 |
| Splice donor | 86.8 ± 0.6 | 88.2 ± 0.8 | 97.6 ± 0.3 | 97.0 ± 0.2 | 78.0 ± 0.7 | 94.3 ± 2.4 | 94.4 ± 2.6 | 96.4 ± 1.0 | 84.2 ± 0.9 | 97.7 ± 0.2 | 97.1 ± 0.2 |
| Average rank (full comparison) | **5.11** | **3.61** | **4.89** | **8.06** | **8.06** | **14.00** | **12.78** | **11.11** | **10.39** | **1.67** | **6.50** |

## Appendix E. Hyperparameters per NT task

ModernGENA is fine-tuned using a grid search over learning rates {1 ×10^−5^, 3 ×10^−5^, 5 ×10^−5^}, weight decay {1 ×10^−4^, 1 ×10^−3^, 1 ×10^−2^}, and effective batch sizes {32, 64}. For Modern-GENA, a dropout rate of 0.1 is additionally used in the classification layer. GENA is fine-tuned using a grid search over learning rates {1 ×10^−5^, 3 ×10^−5^, 5 ×10^−5^}, weight decay {1 ×10^−4^, 1 ×10^−3^}, and effective batch sizes 32, 64. The selected hyperparameters for each task are reported in Table 5.

**Table 5:**
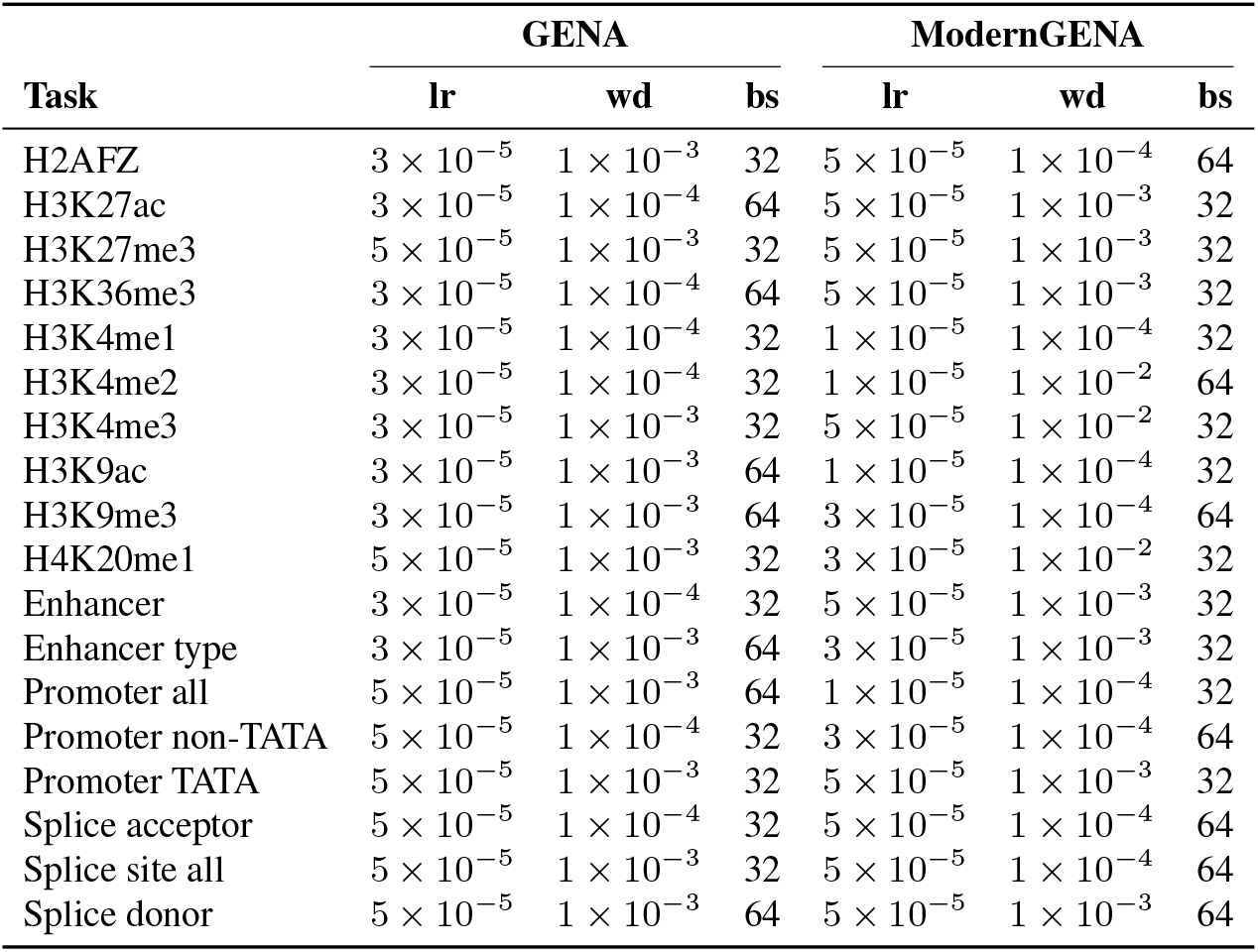
Selected hyperparameters per NT task. lr: learning rate; wd: weight decay; bs: batch size.

| Task | GENA |  |  | ModernGENA |  |  |
| --- | --- | --- | --- | --- | --- | --- |
|  | lr | wd | bs | lr | wd | bs |
| H2AFZ | $3 \times 10^{-5}$ | $1 \times 10^{-3}$ | 32 | $5 \times 10^{-5}$ | $1 \times 10^{-4}$ | 64 |
| H3K27ac | $3 \times 10^{-5}$ | $1 \times 10^{-4}$ | 64 | $5 \times 10^{-5}$ | $1 \times 10^{-3}$ | 32 |
| H3K27me3 | $5 \times 10^{-5}$ | $1 \times 10^{-3}$ | 32 | $5 \times 10^{-5}$ | $1 \times 10^{-3}$ | 32 |
| H3K36me3 | $3 \times 10^{-5}$ | $1 \times 10^{-4}$ | 64 | $5 \times 10^{-5}$ | $1 \times 10^{-3}$ | 32 |
| H3K4me1 | $3 \times 10^{-5}$ | $1 \times 10^{-4}$ | 32 | $1 \times 10^{-5}$ | $1 \times 10^{-4}$ | 32 |
| H3K4me2 | $3 \times 10^{-5}$ | $1 \times 10^{-4}$ | 32 | $1 \times 10^{-5}$ | $1 \times 10^{-2}$ | 64 |
| H3K4me3 | $3 \times 10^{-5}$ | $1 \times 10^{-3}$ | 32 | $5 \times 10^{-5}$ | $1 \times 10^{-2}$ | 32 |
| H3K9ac | $3 \times 10^{-5}$ | $1 \times 10^{-3}$ | 64 | $1 \times 10^{-5}$ | $1 \times 10^{-4}$ | 32 |
| H3K9me3 | $3 \times 10^{-5}$ | $1 \times 10^{-3}$ | 64 | $3 \times 10^{-5}$ | $1 \times 10^{-4}$ | 64 |
| H4K20me1 | $5 \times 10^{-5}$ | $1 \times 10^{-3}$ | 32 | $3 \times 10^{-5}$ | $1 \times 10^{-2}$ | 32 |
| Enhancer | $3 \times 10^{-5}$ | $1 \times 10^{-4}$ | 32 | $5 \times 10^{-5}$ | $1 \times 10^{-3}$ | 32 |
| Enhancer type | $3 \times 10^{-5}$ | $1 \times 10^{-3}$ | 64 | $3 \times 10^{-5}$ | $1 \times 10^{-3}$ | 32 |
| Promoter all | $5 \times 10^{-5}$ | $1 \times 10^{-3}$ | 64 | $1 \times 10^{-5}$ | $1 \times 10^{-4}$ | 32 |
| Promoter non-TATA | $5 \times 10^{-5}$ | $1 \times 10^{-4}$ | 32 | $3 \times 10^{-5}$ | $1 \times 10^{-4}$ | 64 |
| Promoter TATA | $5 \times 10^{-5}$ | $1 \times 10^{-3}$ | 32 | $5 \times 10^{-5}$ | $1 \times 10^{-3}$ | 32 |
| Splice acceptor | $5 \times 10^{-5}$ | $1 \times 10^{-4}$ | 32 | $5 \times 10^{-5}$ | $1 \times 10^{-4}$ | 64 |
| Splice site all | $5 \times 10^{-5}$ | $1 \times 10^{-3}$ | 32 | $5 \times 10^{-5}$ | $1 \times 10^{-4}$ | 64 |
| Splice donor | $5 \times 10^{-5}$ | $1 \times 10^{-3}$ | 64 | $5 \times 10^{-5}$ | $1 \times 10^{-3}$ | 64 |

## Appendix F. Genomic assemblies for multispecies pretraining

**Table 6:**
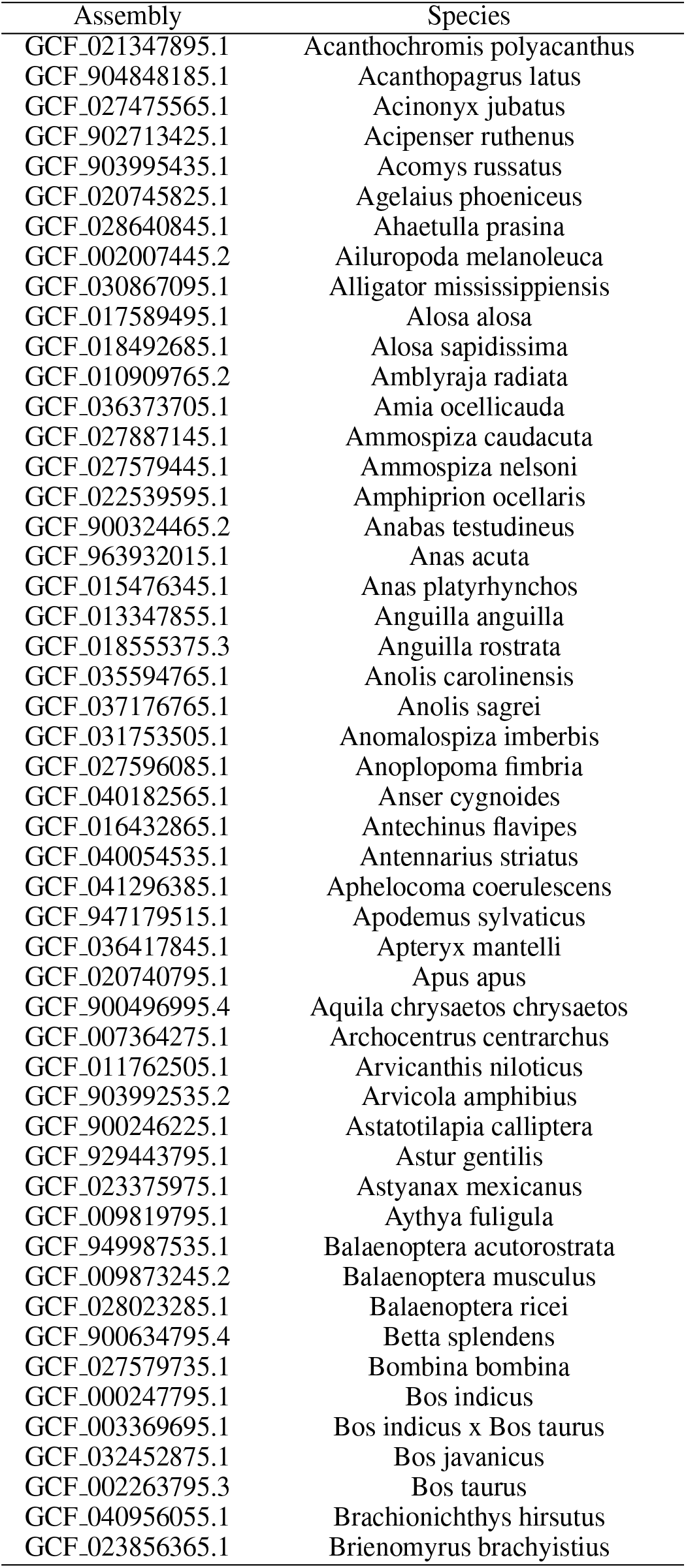
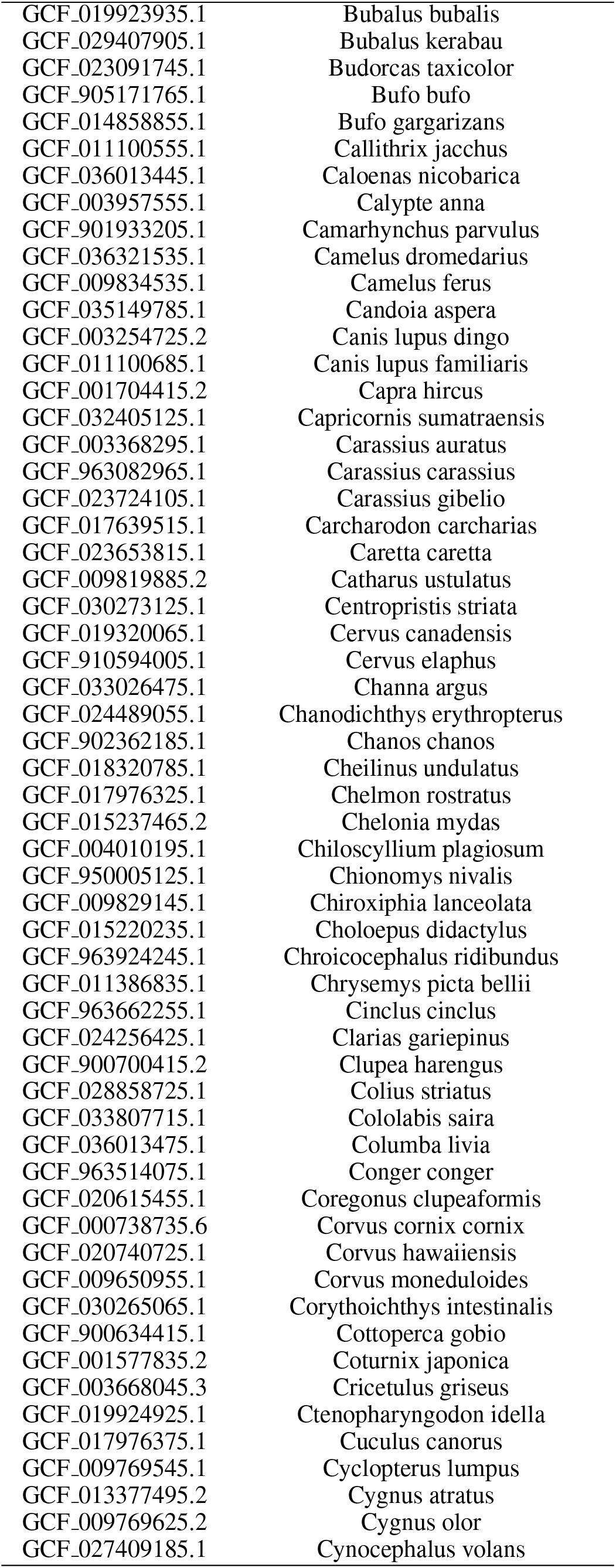
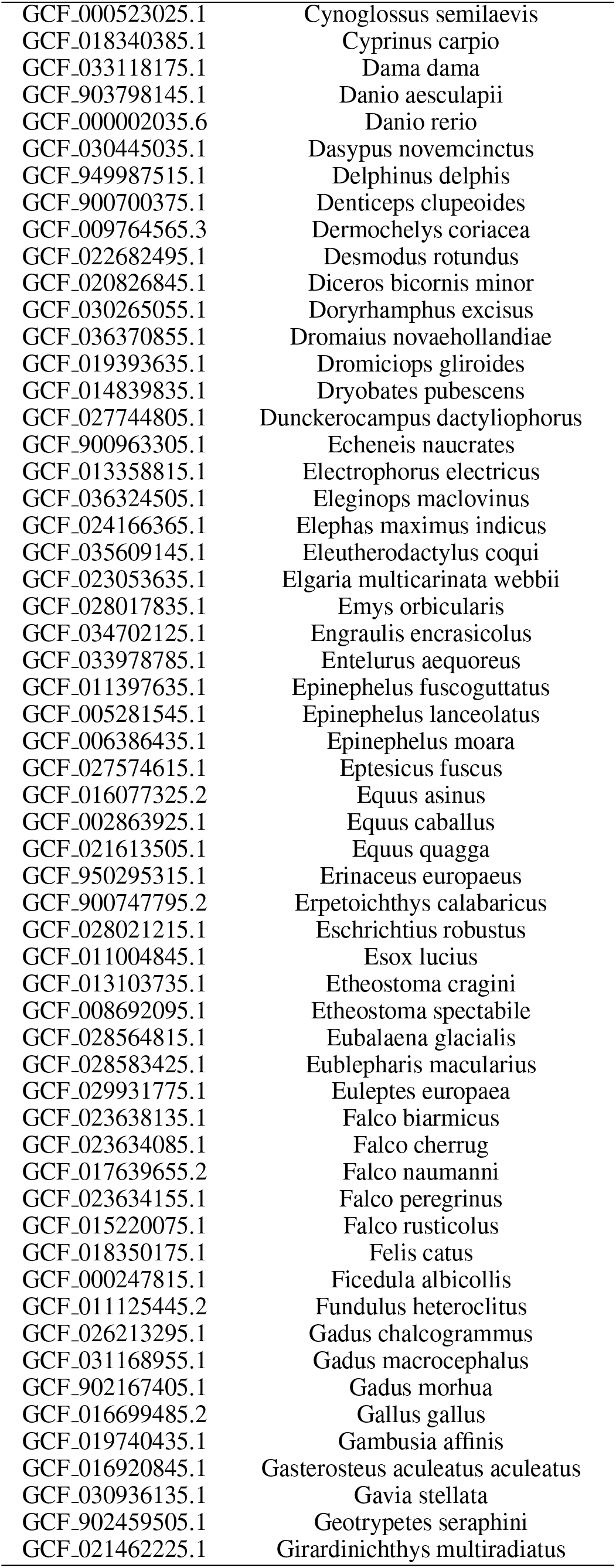
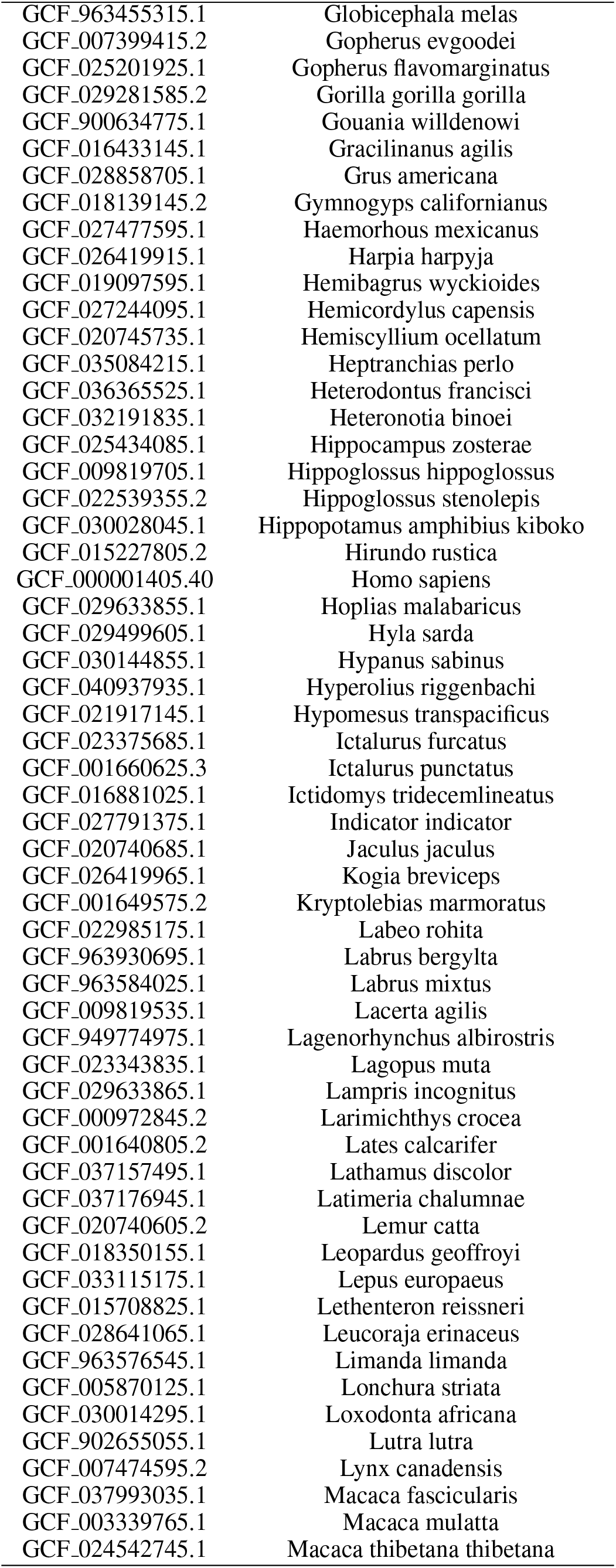
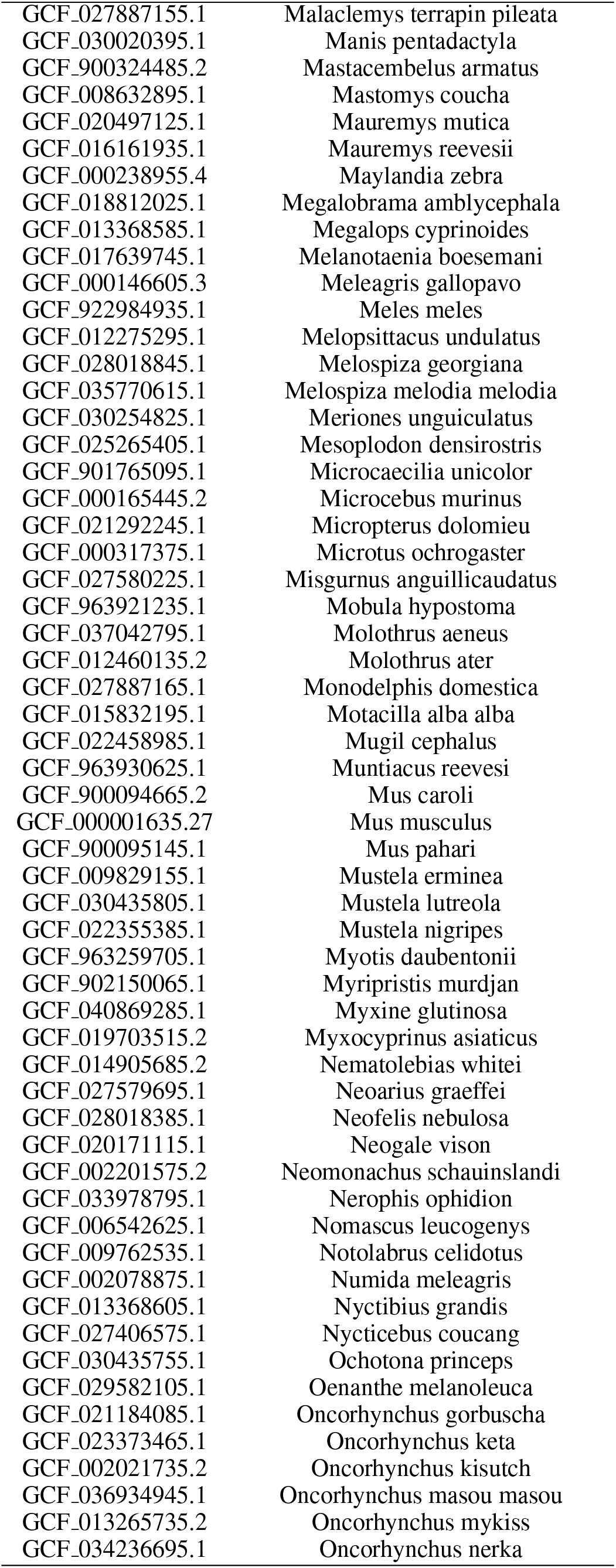
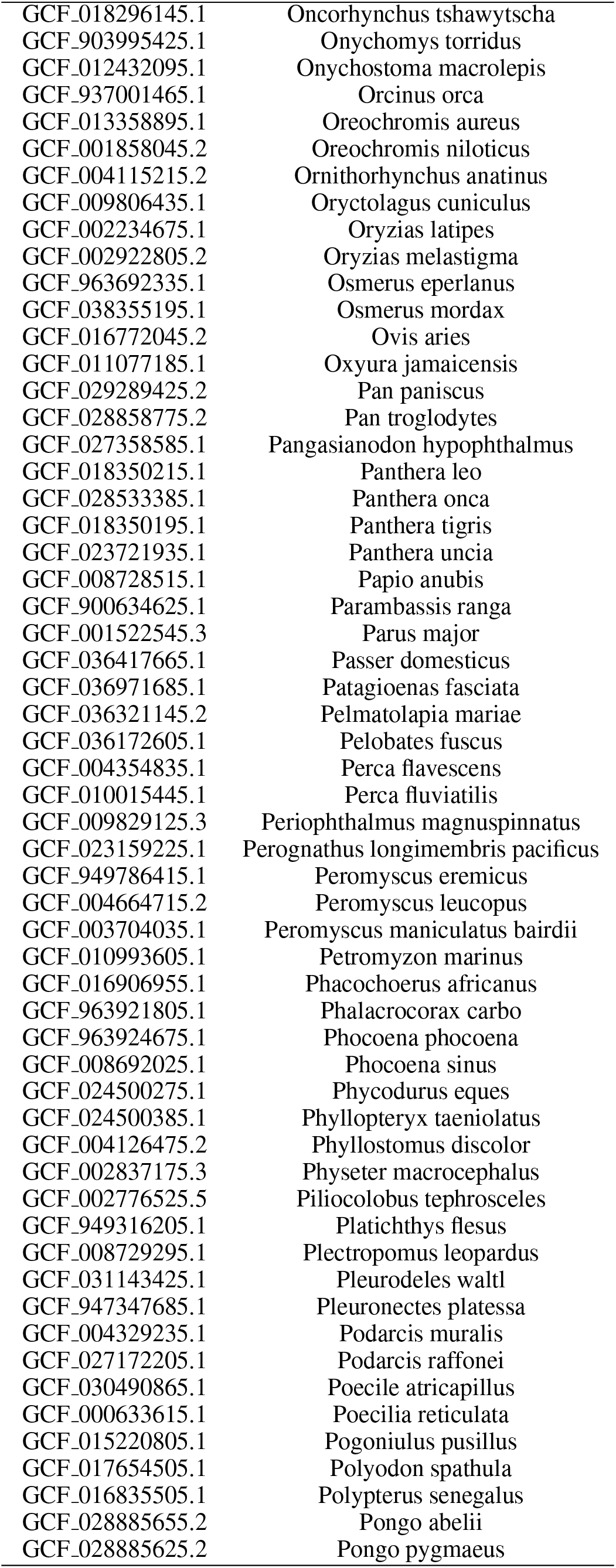
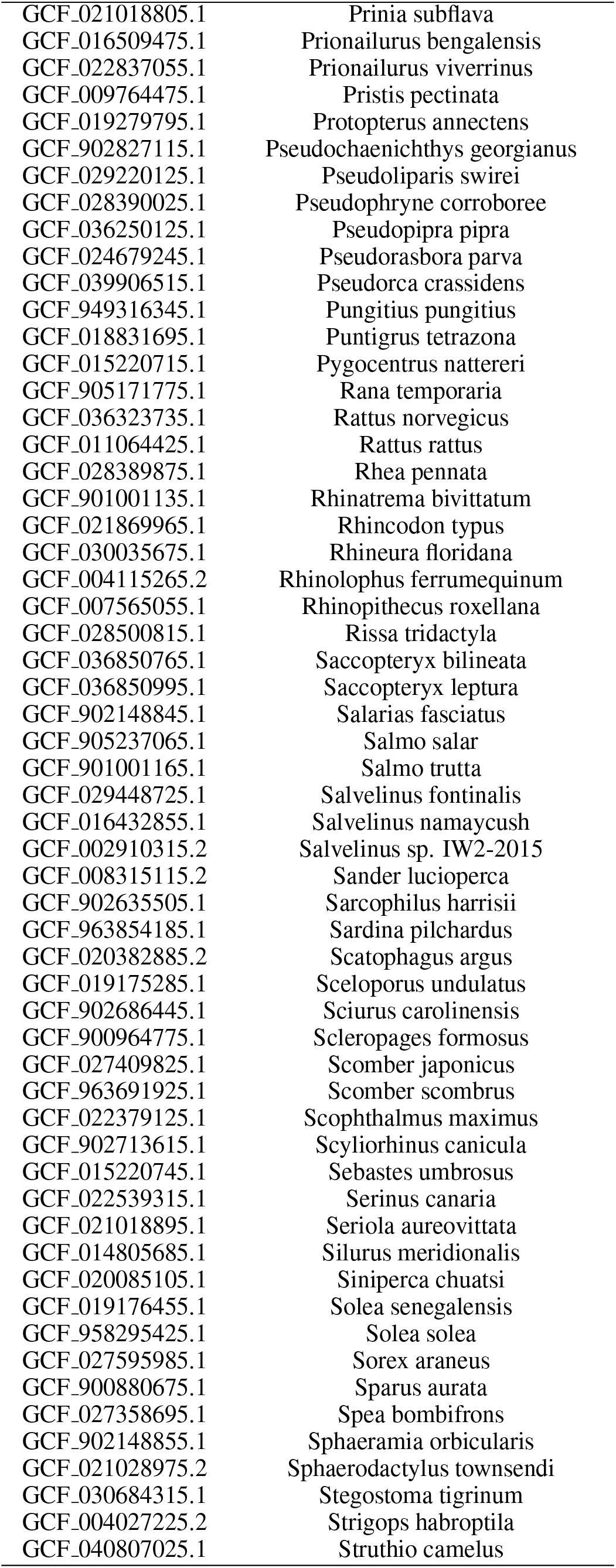
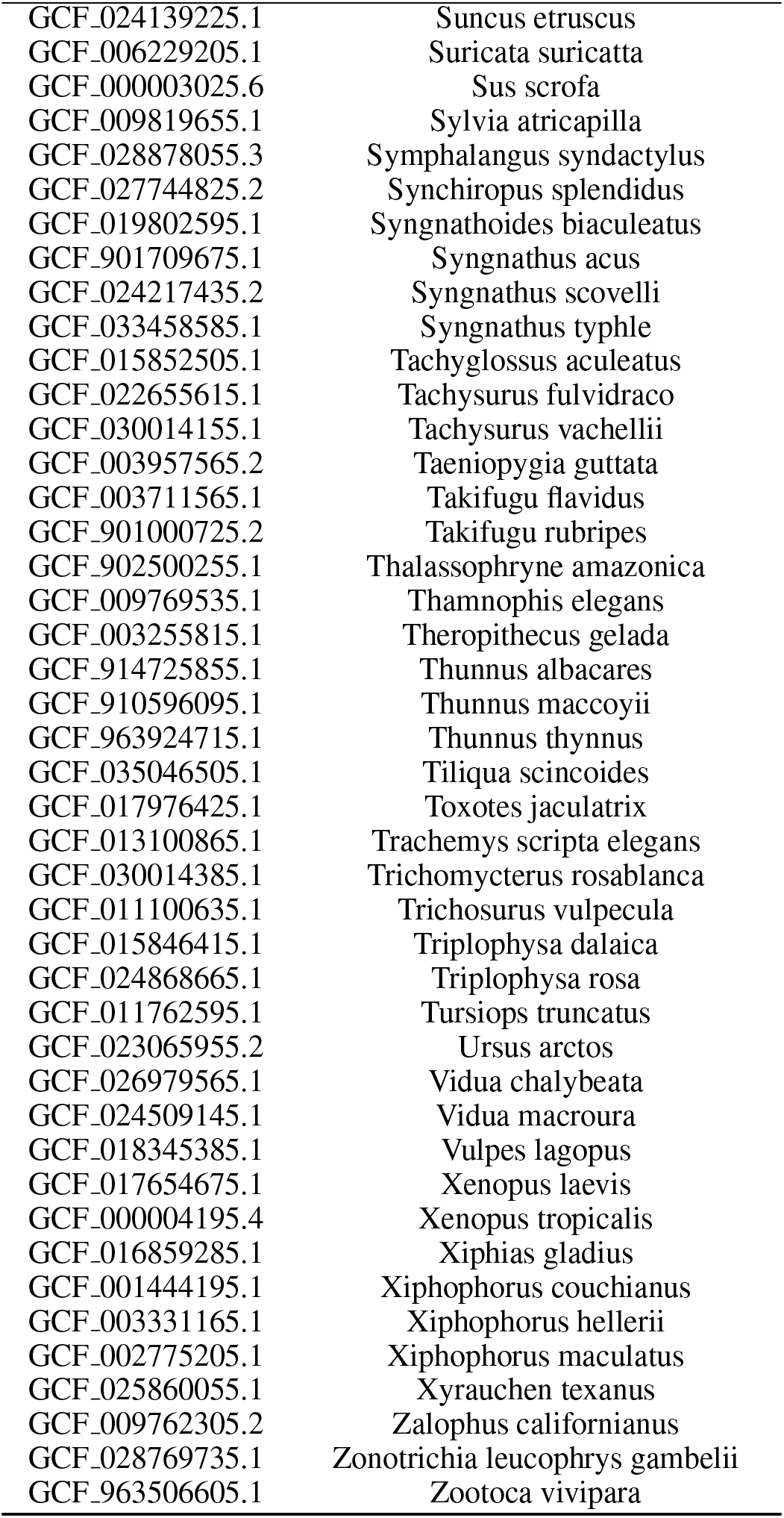
**List of genomic assemblies** used to create the multispecies pretraining dataset. Assembly accessions correspond to the genome and annotation identifiers processed by the NCBI Eukaryotic Genome Annotation Pipeline.

